# Path analysis of intra-metastatic hypoxia in breast cancer

**DOI:** 10.1101/2022.11.03.515032

**Authors:** Sergio Rey-Keim, Luana Schito

## Abstract

Hypoxia (low O_2_) signals into the nucleus of cancer cells through hypoxia-inducible factor (HIF)-1α-dependent transcription triggering proliferative, metabolic and vascular adaptations linked to therapy resistance and mortality due to overt metastasis. In contrast with the wealth of molecular data on primary intra-tumoral hypoxia, there is a dearth of statistical modelling studies addressing the mechanisms of intra-metastatic hypoxia. In this study, we used path analysis to model intra-metastatic hypoxia (Hx), HIF-1α expression and microvascular area (MVA) as functions of metastatic cross-sectional area (MCSA) in an advanced mouse breast cancer model; in this context, we tested the effect of conventional, maximum-tolerated dose (MTD) or low-dose metronomic (LDM) chemotherapy. Iterative analysis of 34 non-isomorphic paths yielded four well-fitting, configuration-invariant models [*χ*^2(6,171)^ ≥ 6.12; *P* ≥ 0.328; CFI ≥ 0.998; RMSEA ≤ 0.07]. All four models contained HIF-1α as a mediating variable within the MCSA↔Hx↔MVA path, as well as significant Hx↔MVA interactions. LDM disrupted the HIF-1α↔MCSA→Hx and HIF-1α↔MVA paths; furthermore, all LDM and MTD combinations impaired Hx↔HIF-1α. These results confirmed well-established hypoxic interactions, whilst uncovering possible differential effects of chemotherapeutic modalities upon metastatic size, hypoxia, HIF-1α and vascularisation. Our data indicate that MVA can act as a downstream readout, rather than an adaptive angiogenic mechanism alleviating intra-metastatic hypoxia. Moreover, well-fitting path models locate HIF-1α activity either upstream or downstream of hypoxia, thereby allowing us to posit the existence of bi-directional feedback loops driving vascularisation and growth in metastatic tumours, of relevance for targeted therapies.

## Background

Hypoxia (low O_2_) is a ubiquitous microenvironmental feature of solid cancers, associated with progression and poor survival due to overt metastatic dissemination, wherein cancer cells (CCs) escape the primary tumour *via* angiogenic vessels (Schito & Semenza, 2016; Rey *et al*, 2017; Schito, 2019). Intra-tumoral hypoxia drives angiogenesis and metastasis through the activation of a transcriptional program dependent upon hypoxia-inducible factor (HIF)-1α and -2α [heretofore referred to as HIFα] (Schito & Semenza, 2016). Importantly, HIFα are thought to activate ≈1-10% of the human transcriptome (Manalo *et al*, 2005; Schito & Semenza, 2016), endowing CCs with the ability to co-opt a panoply of metabolic, angiogenic, migratory and immunoevasive adaptations (Lou *et al*, 2011; Schito & Semenza, 2016; Rey *et al*, 2017) allowing CCs to escape the primary tumour and metastasize onto distant sites, a progression event that is responsible for ≈67% of cancer deaths (Dillekås *et al*, 2019).

Of relevance to the present study, breast cancer is the most common malignancy in women, where recurrence and metastasis have been linked to hypoxia and HIFα induction (Bos *et al*, 2003; Lou *et al*, 2011; Gilkes & Semenza, 2013; Schito & Rey, 2017). Despite considerable advances in the understanding of molecular mechanisms underlying hypoxic metastasis in breast cancer, the emergence of high-throughput data at the genomic, transcriptomic and proteomic levels has uncovered additional, layered levels of complexity to be reconciled with well-established, detailed knowledge on specific transduction pathways (Winslow *et al*, 2012; Tiwary, 2020). From this perspective, the use of statistical and mathematical approaches allow a complementary point-of-view to deconvolve the effect of hypoxia upon metastatic progression. Herein, we utilized an iterative modelling approach *via* path analysis, a statistical tool originally developed for population genetics (Wright, 1934), allowing us test directional interactions among variables in order to identify statistical causal relationships and novel working hypotheses. Path analysis implies the specification of pathways as a system of multiple regression equations configured as a structural model, allowing us to test direct and indirect (mediated) effects, the calculation of overall goodness-of-fit measures and the modification of path configurations in order to optimize model fit (Bollen, 1989; Kline, 2005).

In this study, we modelled the correlation among tumour size [metastatic cross-sectional area (MCSA)], intra-metastatic hypoxia [Hx], HIF-1α expression and microvascular area [(MVA], an index of angiogenesis) in the context of a previously published dataset wherein we tested the effects of conventional, maximum-tolerated dose (MTD) chemotherapy against continuous, low-dose metronomic (LDM) regimens upon HIFα levels in a mouse model of advanced breast cancer that spontaneously metastasizes to the lungs (Schito *et al*, 2020). Systematic analysis of path configurations revealed interactions that are in line with our current understanding of hypoxic HIFα signalling; furthermore, we observed statistical evidence of HIFα-independent mechanisms controlling angiogenic responses in hypoxic metastatic nodules. Or data revealed that doublet and singlet MTD and LDM chemotherapies disrupted distinct paths, thereby suggesting divergent mechanisms of action. Furthermore, our data suggest that microvascular density correlates with the severity of intra-metastatic hypoxia and therefore is unsuitable as a predictor for antiangiogenic therapy responses, since it would reflect a poorly perfused metastatic vasculature that, in principle, would make cancer cells inaccessible to most orally and intravenously administered therapies.

## Methods

### Metastatic breast cancer model and chemotherapy administration

Data were gathered from a previous study in a syngeneic metastatic breast cancer model (Schito *et al*, 2020). Briefly, cisplatin-resistant, mammary adenocarcinoma [EMT6-CDDP cells (100,000)] were orthotopically implanted in the inguinal mammary fat pad of fifty immunocompetent female mice (Teicher *et al*, 1990; Shaked *et al*, 2016; Schito *et al*, 2020). Primary breast tumours were resected after 12 days, before initiating adjuvant LDM or MTD chemotherapy; treatment was sustained for ten days whilst monitoring for overt signs of metastatic disease. Cyclophosphamide ([CTX], Baxter) was administered at 20 mg/kg/day PO through the drinking water, whereas capecitabine ([CPB], LC Laboratories) was given at continuous LDM doses of 100 mg/kg/d PO by gavage or as bolus MTD doses of 400 mg/kg/d PO for 4 d, followed by a 17d drug-free break period. Lung tissue was collected at the end of each chemotherapy administration protocol.

### Quantitative immunohistochemistry and morphometry

Tissue processing and antigen retrieval were performed as previously described (Schito *et al*, 2012, 2020); briefly, 5-μm thick lung tissue sections were submitted for immunohistochemistry with primary anti-HIF-1α, HIF-2α, pimonidazole [hypoxia [Hx] marker (Nordsmark *et al*, 2003; Ragnum *et al*, 2015)] or CD31 (endothelial marker [MVA]), as previously described (Schito *et al*, 2020). Microphotographs (≈ 1,200 digital images) were acquired at ×100 magnification (1.3 μm/px) with an IMX533 CMOS camera (Sony Corp., Japan) coupled to a Zeiss Primostar 3 microscope (Carl Zeiss AG, Germany) under fixed conditions for LED illumination, exposure and flatfield correction. Digitally stitched images were subsequently submitted to unsupervised, automated quantitative immunohistochemistry using custom code run within ImageJ (v1.53q, NIH). Metastatic nodules were segmented into individual images, deconvoluted into haematoxylin and diaminobenzidine signals, and processed to determine total HIF-1α^+^, HIF-2α^+^, pimonidazole^+^ and CD31^+^ signals at fixed pixel intensity thresholds in consecutive histological sections. MCSA was calculated in parallel as the average total area in all sections belonging to each metastatic nodule.

### Directed path enumeration

In order to evaluate all possible combinatorial paths among MCSA, Hx, HIF-1α and MVA, we assessed path structures as directed graphs with four vertices, thereby enumerating 34 ‘core’ out of all the 38 possible structural path models considering four vertices (Harary & Palmer, 1973). Only non-isomorphic directed paths were considered.

### Iterative screening of path models

Systematic assessment of global model fit was carried out based on the results of directed path enumeration within a graphical environment (SPSS Amos v24.0.0, IBM Corp., Wexford, MA). Models were tested using a maximum likelihood approach and Monte Carlo bootstrapping permutations (5,000). A non-significant χ^2^ statistic (*P* > 0.05) combined with a χ^2^:*df* ≤ 3 ratio and a root mean square error of approximation (RMSEA) ≤ 0.07, were used as cut-off points to select candidate models for further analysis (Steiger, 2007). When these *criteria* were fulfilled, the model was tested for improved fit by adding covariances in the context of known, experimentally validated correlations known to occur in cancer models and clinical studies of intra-tumoral hypoxia (*i*.*e*., MCSA↔HIF-1α, HIF-1α↔MVA, Hx↔MVA and MCSA↔Hx) (Generali *et al*, 2006; Rey & Semenza, 2010; Schito & Rey, 2020).

### Statistical analysis

#### Data analysis

All statistical analyses, except for the iterative screening of path models were carried out in *R* (v. 4.1.3) using the packages *dplyr* (v. 1.0.8), *lavaan* (v. 0.6-11.1676) (Rosseel, 2012), *lavaan*.*survey* (v. 1.1.3.1), *lavaanPlot* (v. 0.6.2), *semTools* (v. 0.5-5), and *rcompanion* (v. 2.4.15). All data were expressed as means ± SEM unless noted otherwise. Robust estimates were used for all statistical analyses.

#### Data transformations

Calibrated positive areas for MCSA, Hx, HIF-1α and MVA were expressed in μm^2^. In order to account for samples where immunoreactivities were below detection limit, and thus equalled zero, data were transformed with a log_10_[*x*+1] function, followed by normalization with the Tukey’s ladder of powers method (λ interval = 0.003), followed by standardization onto *t*-values. Data were then tested for normality *via* Shapiro-Wilk and Anderson-Darling tests as well as with quantile-quantile (QQ) plots. Next, a *Spearman* ρ correlation matrix was built to explore the direction and strength of relationships in paired comparisons among MCSA, Hx, HIF-1α and MVA. A total of 171 pulmonary metastatic nodules with no missing values were analysed.

#### Final model fit

Path analysis of all four selected models was carried out with the *lavaan* package in *R*, using a Satorra-Bentler scaled test statistic (Satorra & Bentler, 2001). Multi-group analysis was done by grouping samples according to six distinct treatment groups: vehicle control, low-dose metronomic capecitabine singlet (LDM^CTX^), maximum-tolerated dose capecitabine singlet (MTD^CPB^), low-dose metronomic cyclophosphamide singlet (LDM^CTX^), LDM doublet cyclophosphamide + capecitabine (LDM^CTX^ + LDM^CPB^) and doublet low-dose metronomic cyclophosphamide + maximum-tolerated dose capecitabine (LDM^CTX^ + MTD^CPB^). Robust measures of fit were calculated for all models: *Akaike* information criterion (AIC), Bayesian information criterion (BIC), comparative fit index (CFI), scaled *χ*^*2*^, root mean square error of approximation (RMSEA), standardized root mean square residual (SRMR) and *Tucker-Lewis* index (TLI). The statistical significance of standardized loadings for each path within a model was tested in a group-by-group basis using a *t*-statistic (H_1_: loadings ≠ 0).

#### Measurement invariance testing

For each converging model with acceptable measures of fit (configural invariance), we sequentially assessed measurement invariance by testing the unconstrained model against two nested models: *first*, a model with fixed loadings per path were equal across groups (metric/weak invariance) and *second*, a model wherein all loadings per path were equal as well as all intercepts (scalar/strong invariance). ANOVA testing was performed using the Satorra-Bentler scaled χ^2^ statistic as implemented within the *lavaan R* package (Satorra & Bentler, 2010; Rosseel, 2012).

## Results

### Intra-metastatic hypoxia, vascularisation and tumour size

Morphometric analysis of individual metastatic nodules from EMT6-CDDP primary breast tumours showed that nodular area, HIF-1α, intra-metastatic hypoxia, and vascularisation signals were not normally distributed (*P*< 2.2×10^−16^ by both Shapiro-Wilk and Anderson-Darling tests); rather, they all followed right-tailed lognormal distributions. Logarithmic transformations, followed by the Tukey’s ladder of power function attained normalisation in QQ plots (*P* ≥ 0.596, Fig. 1A). Remarkably, analysis of cross-correlations among MCSA, Hx, HIF-1α and MVA consistently showed positive *Spearman* rank correlation coefficients [0.50 ≤ ρ ≤ 0.88; *P* ≤ 1.41×10^−7^; *n* = 171; Fig. 1B]. Importantly, it should be noted that lung metastatic nodules from EMT6-CDDP breast tumours did not express HIF-2α (*data not shown*), thus making HIFα signalling reliant exclusively upon HIF-1α.

**Figure 1.**
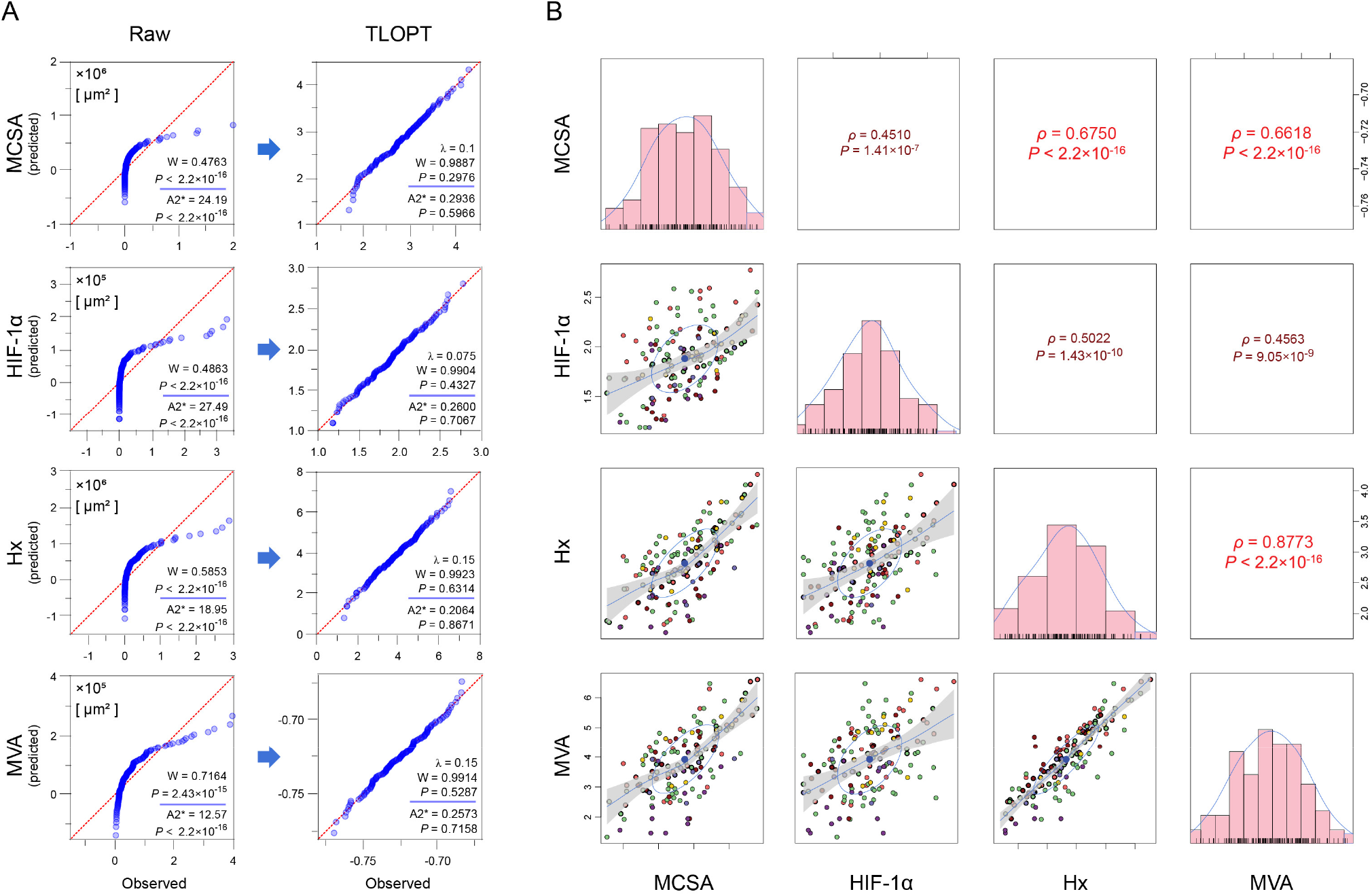
Data distribution and correlations among nodule size, hypoxia, HIF-1α and vascularisation in 171 pulmonary breast cancer metastases. **A**, Quantile-quantile plots showing the difference between observed and predicted gaussian data before (Raw, *left column*) and after normalization with a *Tukey’s ladder of powers transformation* (TLOPT, *right column*). Normality was assessed with *Shapiro-Wilk* and *Anderson-Darling* tests before and after data transformation. **B**, *Spearman* correlation matrix and scatterplots showing the relationship among measured variables. Graphs along the diagonal show data distributions after TLOPT. *P*, significance by *F*-tests (slope ≠ 0). *Blue*, best-fitting spline regressions; *gray*, 95% confidence interval of the regression line. *A2\**, Anderson-Darling statistic; *HIF-1α*, hypoxia-inducible factor-1α; *Hx*, hypoxia; *MCSA*, metastatic cross-sectional area; *MVA*, metastatic vascular area; *W*, Shapiro-Wilk statistic; *λ*, stepwise interval for TLOP transformation; *ρ*, Spearman’s *rho* rank correlation coefficient.

In order to explore whether the statistical interactions among metastatic size, hypoxia, HIF-1α expression and vascularisation contained bi-directional interaction or feedbacks, we estimated four distinct general linear models (GLMs, henceforth referred to as L1, L2, L3 and L4), correlating MCSA, Hx, HIF-1α and MVA with mouse ID as a fixed effect whilst stratifying by chemotherapy group defined as follows:

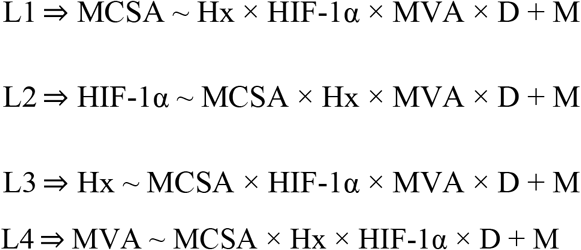

#### Wherein

*D* is the drug regimen [vehicle, LDM^CTX^, MTD^CPB^, LDM^CTX^, LDM^CTX^ + LDM^CPB^ or LDM^CTX^ + MTD^CPB^] and *M* represents individual mouse IDs.

Interestingly, the effect of hypoxia was statistically significant in all GLMs where it acted as a predictor (L1 [for MCSA], L3 [for Hx] and L4 [for MVA]; *P* < 2×10^−16^) whilst interacting with HIF-1α in L1 (*P* = 0.041). Metastatic size exerted a strong predictive effect in L2, L3 and L4, wherein it acted as a predictor upon HIF-1α, hypoxia and vascularisation (*P* ≤ 8.96 × 10^−5^). Further, HIF-1α levels acted as a predictor of metastatic size (*P* = 0.002) and hypoxia (*P* < 2×10^−16^) in models L1 and L3; however, HIF-1α expression did not modify vascularisation in L4 (*P* = 0.079). Metastatic vascular area was likewise associated with metastatic size (L1: *P* = 0.042), HIF-1α levels (L2: *P* = 0.031) and hypoxia (L3: *P* < 2×10^−16^). Importantly, both the effects of chemotherapy and individual differences among experiments were significant in all models (*P* < 0.01). Taken together, integration of all four GLMs suggests that, except for the interaction between hypoxia and HIF-1α, the modelled variables are orthogonal and thus feasible to be modelled by including feedback loops.

### Modelling of intra-metastatic hypoxia

Directed path enumeration resulted in 34 acyclical, non-isomorphic models that we classified according to their geometry among six types. That is, ‘diamond’, ‘paw’, ‘trident’, ‘4-path’, ‘square’ and ‘pyramid’ (Fig. 2A-F). Sequential testing of model fit in all configurations indicated that most convergent models were located in the path categories ‘diamond’ (Fig. 2A) and ‘square’ (Fig. 2E). Based upon GLM results, we tested nested variations of the converging models wherein we replaced the different paths with covariates as appropriate; this iterative process yielded four final models, herein termed M1 to M4 (Fig. 3A-D). Of note, models M1 and M2 featured the canonical mode of regulation of HIF-1α transactivation (HIF-1α←Hx) whilst M3 and M4 reversed the path (Hx←HIF-1α). All tested measurements of model fit showed a concordant ranked order from best to worst fit (M1>M4>M2>M3); that is, non-significant chi-squared scaled fits [*χ*^2(6,171)^ ≥ 6.12; *P* ≥ 0.328] as well as meeting conventionally accepted thresholds for alternative fit indexes [*i*.*e*., CFI > 0.95 ; TLI > 0.95 and RMSEA < 0.06 except for M3, wherein RMSEA = 0.073 (Table 1)] (Bentler, 1990; Kenny *et al*, 2015). A non-significant overall *χ*^2^ indicates that there is no difference between the data matrix predicted by each model and the measured data matrix. *In lieu* of recent data suggesting that RMSEA might be overestimated in models with small degrees of freedom (Kenny *et al*, 2015; Shi *et al*, 2021), we decided to include M3 among the final models, considering that *χ*^2^, CFI and TLI indexes were favourable. These results indicate that the overall factor structure for all models is similar in all five chemotherapeutic regimens as well as in the vehicle group (i.e., the models satisfy configural invariance).

**Figure 2.**
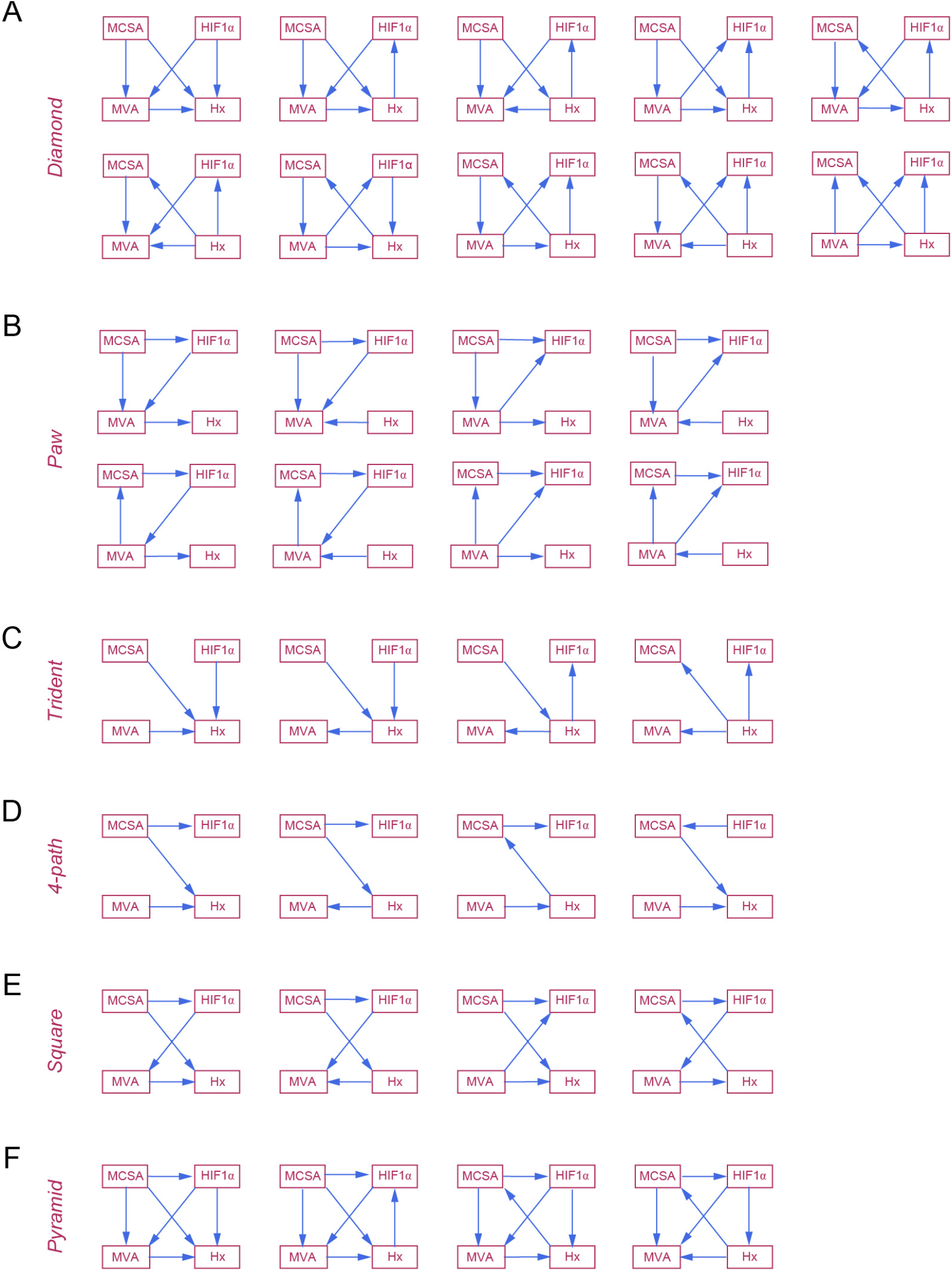
Graphical representation of all 34 acyclical candidate directed path models. According to their configuration, candidate path models were classified among six categories: **A**, *diamond*; **B**, *paw*; **C**, *trident*; **D**, *4-path* (linear); **E**, *square* and **F**, *pyramid*. Only non-isomorphic paths were considered. These diagrams do not include nested models wherein interactions were replaced with covariates. *HIF-1α*, hypoxia-inducible factor-1α; *Hx*, hypoxia; *MCSA*, metastatic cross-sectional area; *MVA*, metastatic vascular area.

**Figure 3.**
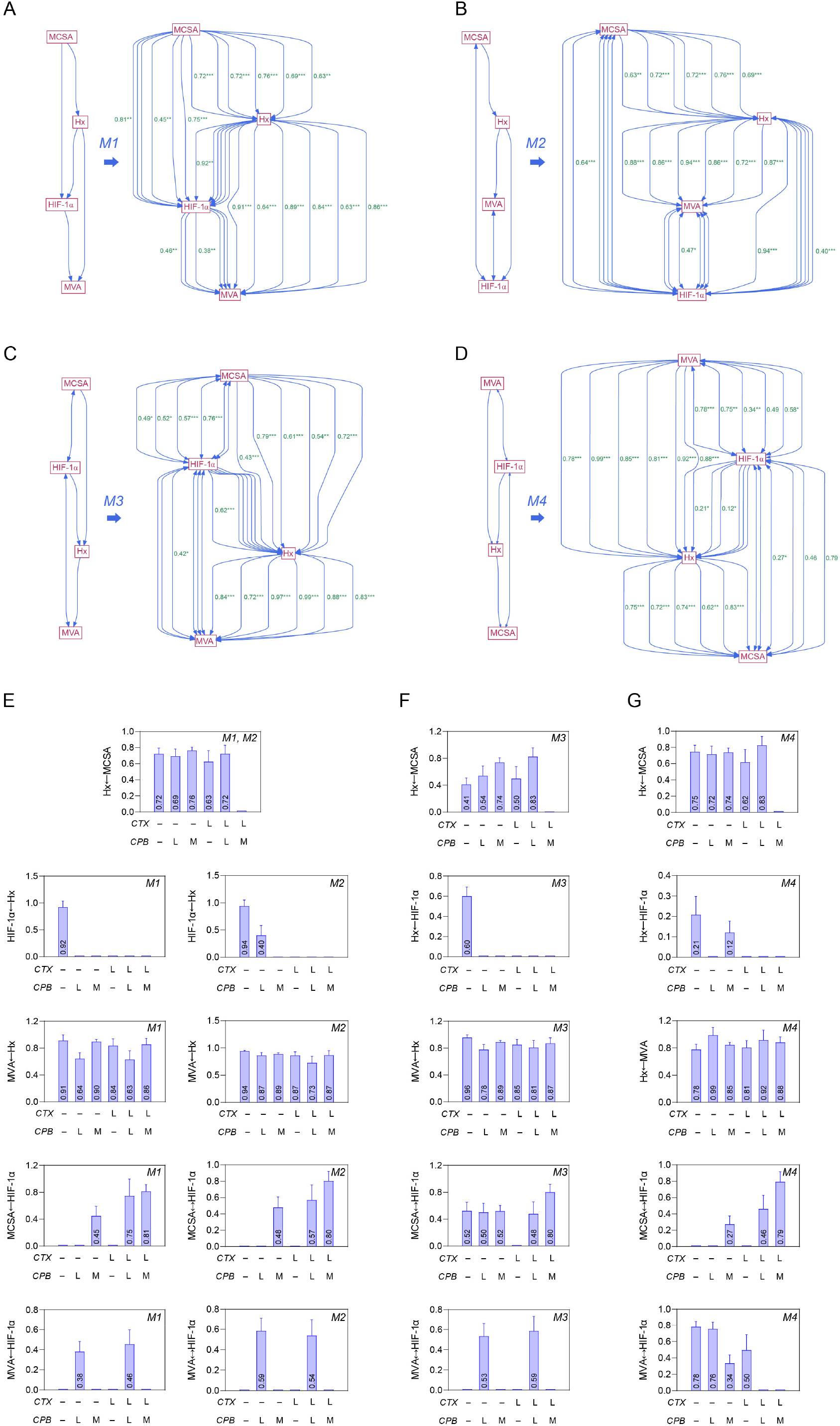
Effect of low-dose metronomic and conventional maximum-tolerated dose chemotherapy upon standardized loadings in all four final path models. **A-D**, Diagrams illustrating the overall configuration (*left*) and individual loadings (*right*) for all six treatment groups using multi-group path analysis. Final models were designated as M1 (A), M2 (B), M3 (C) and M4 (D). Statistical significance of standardized path loadings was tested with a *t*-statistic (loading ≠ 0); ***, *P*< 0.001; **, *P*< 0.01 and *, *P*< 0.05. **E-G**, Individual standardized path loadings for models M1 and M2 (**E**), M3 (**F**) and M4 (**G**). Only statistically significant loadings are plotted as means ± SEM. All non-significant standardized loadings are assumed to be equal to zero. *CTX*, cyclophosphamide; *CPB*, capecitabine; *HIF-1α*, hypoxia-inducible factor-1α; *Hx*, hypoxia; *L*, low-dose metronomic chemotherapy; *M*, maximum-tolerated dose chemotherapy; *MCSA*, metastatic cross-sectional area; *MVA*, metastatic vascular area. *Double arrows* indicate covariances, whereas *single arrows* indicate unidirectional paths. Individual path features are depicted in *green*.

**Table 1:**
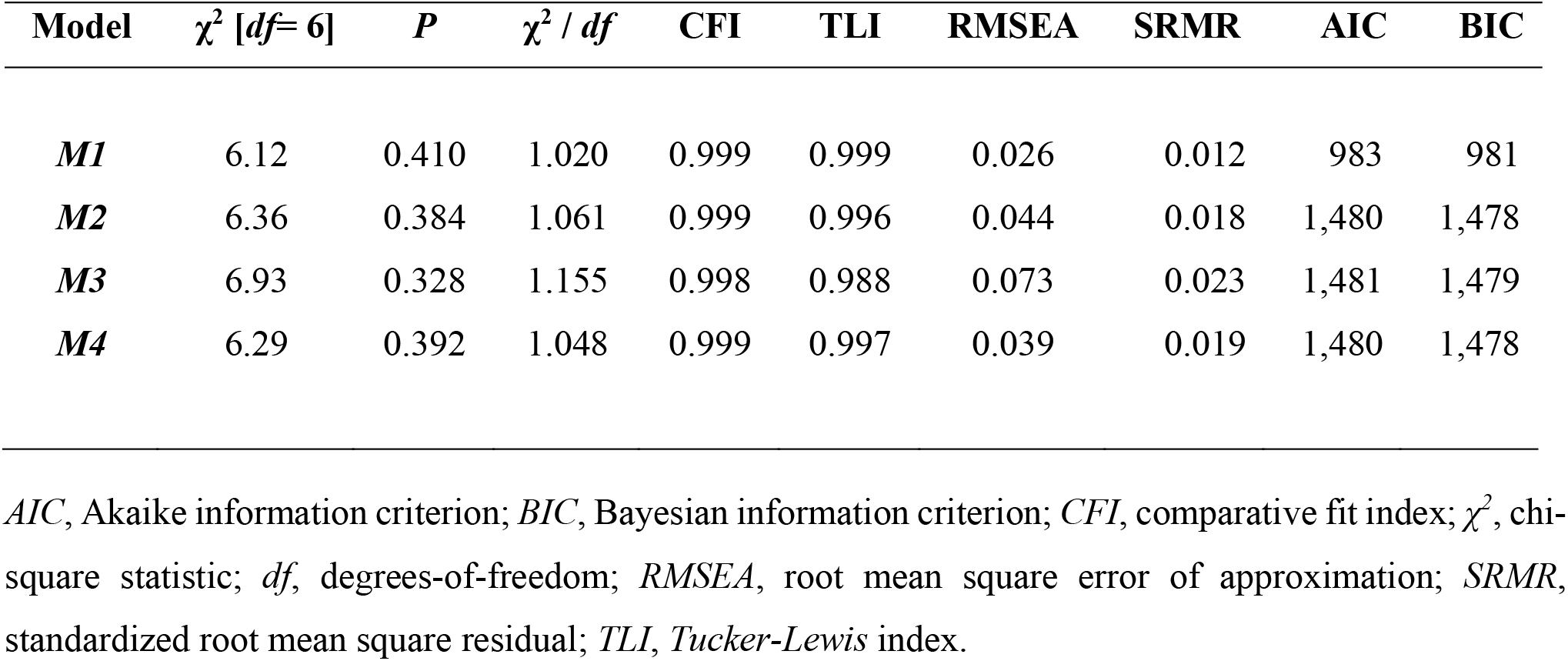
Measures of fit for final path models of intra-metastatic hypoxia.

| **Model** | **$\chi^2$ [*df*= 6]** | ***P*** | **$\chi^2$ / *df*** | **CFI** | **TLI** | **RMSEA** | **SRMR** | **AIC** | **BIC** |
| --- | --- | --- | --- | --- | --- | --- | --- | --- | --- |
| ***M1*** | 6.12 | 0.410 | 1.020 | 0.999 | 0.999 | 0.026 | 0.012 | 983 | 981 |
| ***M2*** | 6.36 | 0.384 | 1.061 | 0.999 | 0.996 | 0.044 | 0.018 | 1,480 | 1,478 |
| ***M3*** | 6.93 | 0.328 | 1.155 | 0.998 | 0.988 | 0.073 | 0.023 | 1,481 | 1,479 |
| ***M4*** | 6.29 | 0.392 | 1.048 | 0.999 | 0.997 | 0.039 | 0.019 | 1,480 | 1,478 |

*AIC*, Akaike information criterion; *BIC*, Bayesian information criterion; *CFI*, comparative fit index; $\chi^2$ , chi-square statistic; *df*, degrees-of-freedom; *RMSEA*, root mean square error of approximation; *SRMR*, standardized root mean square residual; *TLI*, *Tucker-Lewis* index.

We next tested for metric (weak) and scalar (strong) invariance across treatment groups for M1, M2, M3 and M4 by comparing each unconstrained model against a fixed-loadings or a fixed loadings/intercepts in a nested configuration. Interestingly, metric and scalar invariance tests were statistically significant for all four models (*P* < 0.01), indicating that both LDM and MTD chemotherapy regimens exert non-linear effects on the path coefficients and intercepts, nonetheless within structurally equivalent models (Table 2). These results prompted us to assess the effect of chemotherapy regimens in terms of significance by *t*-statistics, nonetheless precluding us from assessing differences among the magnitude of loadings *per se*.

**Table 2:** Invariance testing of final path models of intra-metastatic hypoxia.

| Model | Invariance type ($H_0$) | AIC | BIC | $\chi^2$ [df] | $\Delta\chi^2$ [df] | CFI [$\Delta$] | $P$ |
| --- | --- | --- | --- | --- | --- | --- | --- |
| **M1** | Configural | 983 | 981 | 6.12 [6] | – | 1.000 [0.000] | 0.410 |
|  | Metric | 985 | 985 | 63.10 [31] | 56.98 [25] | 0.937 [0.063] | 0.00023 |
| | Scalar | 1,528 | 1,527 | 199.27 [55] | 136.17 [24] | 0.767 [0.170] | $<2.2 \times 10^{-16}$ |
| **M2** | Configural | 1,480 | 1,478 | 6.36 [6] | – | 0.999 [0.000] | 0.384 |
|  | Metric | 1,480 | 1,479 | 56.18 [31] | 49.82 [25] | 0.947 [0.052] | 0.00256 |
| | Scalar | 1,527 | 1,526 | 189.75 [55] | 133.57 [24] | 0.773 [0.174] | $<2.2 \times 10^{-16}$ |
| **M3** | Configural | 1,481 | 1,479 | 6.92 [6] | – | 0.998 [0.000] | 0.328 |
|  | Metric | 1,483 | 1,482 | 60.05 [31] | 53.13 [25] | 0.939 [0.056] | 0.00088 |
| | Scalar | 1,527 | 1,527 | 190.54 [55] | 130.49 [24] | 0.772 [0.148] | $<2.2 \times 10^{-16}$ |
| **M4** | Configural | 1,480 | 1,478 | 6.29 [6] | – | 0.999 [0.000] | 0.392 |
|  | Metric | 1,473 | 1,471 | 51.03 [31] | 44.74 [25] | 0.960 [0.039] | 0.00949 |
| | Scalar | 1,517 | 1,517 | 182.17 [55] | 131.14 [24] | 0.791 [0.169] | $<2.2 \times 10^{-16}$ |

*AIC*, Akaike information criterion; *BIC*, Bayesian information criterion; $\chi^2$ , chi-square statistic; *CFI*, comparative fit index [ $\Delta$ = delta CFI]; *df*, degrees-of-freedom; $\Delta\chi^2$ , delta chi-square. *P*, *p*-value (chi-square difference test); $n = 171$ .

### Effect of chemotherapy upon intra-metastatic hypoxia model loadings

After demonstrating configural invariance in all four models, we tested for differential effects of chemotherapy regimens upon intra-metastatic hypoxia by examining the statistical significance of path loadings on each therapy group. Multi-group path analysis was carried out to determine individual loadings in each path for models M1, M2, M3 and M4. Significance was determined by calculating *t*-statistics to determine whether loadings were different from zero.

The effect of nodule size upon hypoxia (Hx←MCSA) was significant in all models, except for the doublet LDM^CTX^ + MTD^CPB^ experimental group (Fig. 3A-D and 3E,F,G [*top row*]). Importantly, HIF-1α acted as a mediating variable between nodule size and hypoxia for all models in the vehicle-treated group, an effect that was abolished by both singlet and doublet chemotherapy modalities, independent of whether administration occurred in an MTD or LDM context (Fig. 3A-D and 3E, F, G [*second row*]); by contrast, loadings correlating metastatic vascularisation with hypoxia (MCSA↔/←HIF-1α) were significant and remained unaffected across models and chemotherapeutic regimens (Fig. 3A-D and 3E,F,G [*third row*]), whereas the interaction between metastatic size and HIF-1α was less consistent, however blocked by LDM^CTX^ in all four models (Fig. 3A-D and 3E,F,G [*fourth row*]). Interestingly, no clear pattern emerged when examining the loadings between metastatic vascularisation and HIF-1α (MVA↔/←HIF-1α; Fig. 3A-D and 3E,F,G [*fifth row*]).

Integration of M1, M2, M3 and M4 (Fig. 4A) revealed a shared model structure, wherein metastatic size was upstream of intra-metastatic hypoxia with HIF-1α as a mediating variable. Notably, hypoxia strongly interacted with metastatic vascularisation, which in turn co-varied with HIF-1α. Importantly, the models hereby analysed suggest that hypoxia signalling does not exclusively occur through Hx→HIF-1α. Instead, our data provides evidence of ‘retrograde’ effects of HIF-1α upon intra-metastatic hypoxia that can be interpreted as the presence of feedback loops at the Hx↔HIF-1α path (Fig. 4A). We speculate that the observed strong interactions among Hx←MCSA and Hx↔MVA are hierarchically overimposed upon HIF-1α↔Hx, HIF-1↔MVA and MCSA←HIF-1α, thereby representing statistical evidence for HIF-1α-independent mechanisms controlling metastatic growth and vascularisation. On the other hand, analysis of the global effect of chemotherapy regimens upon path loadings showed consistent loss-of-correlation of the Hx↔HIF-1α path; by contrast, no chemotherapy combination had a disruptive effect upon the Hx↔MVA interaction (Fig. 4B). Interestingly, different LDM modalities disrupted (or activated) specific pathways; (*i*.*e*., doublet LDM^CTX^ + LDM^CPB^ blocked Hx←MCSA whilst ‘activating’ HIF-1↔MVA, whereas LDM^CTX^ singlet therapy uniquely interfered with the Hx←MCSA interaction). These results suggest that LDM chemotherapy might exert its effects upon intra-metastatic hypoxia by modifying the interaction between metastatic size and hypoxia, whilst having an indirect effect upon HIF-1α transcriptional activity, rather than by inhibiting the transcriptional upregulation of angiogenic factors underlying MVA.

**Figure 4.**
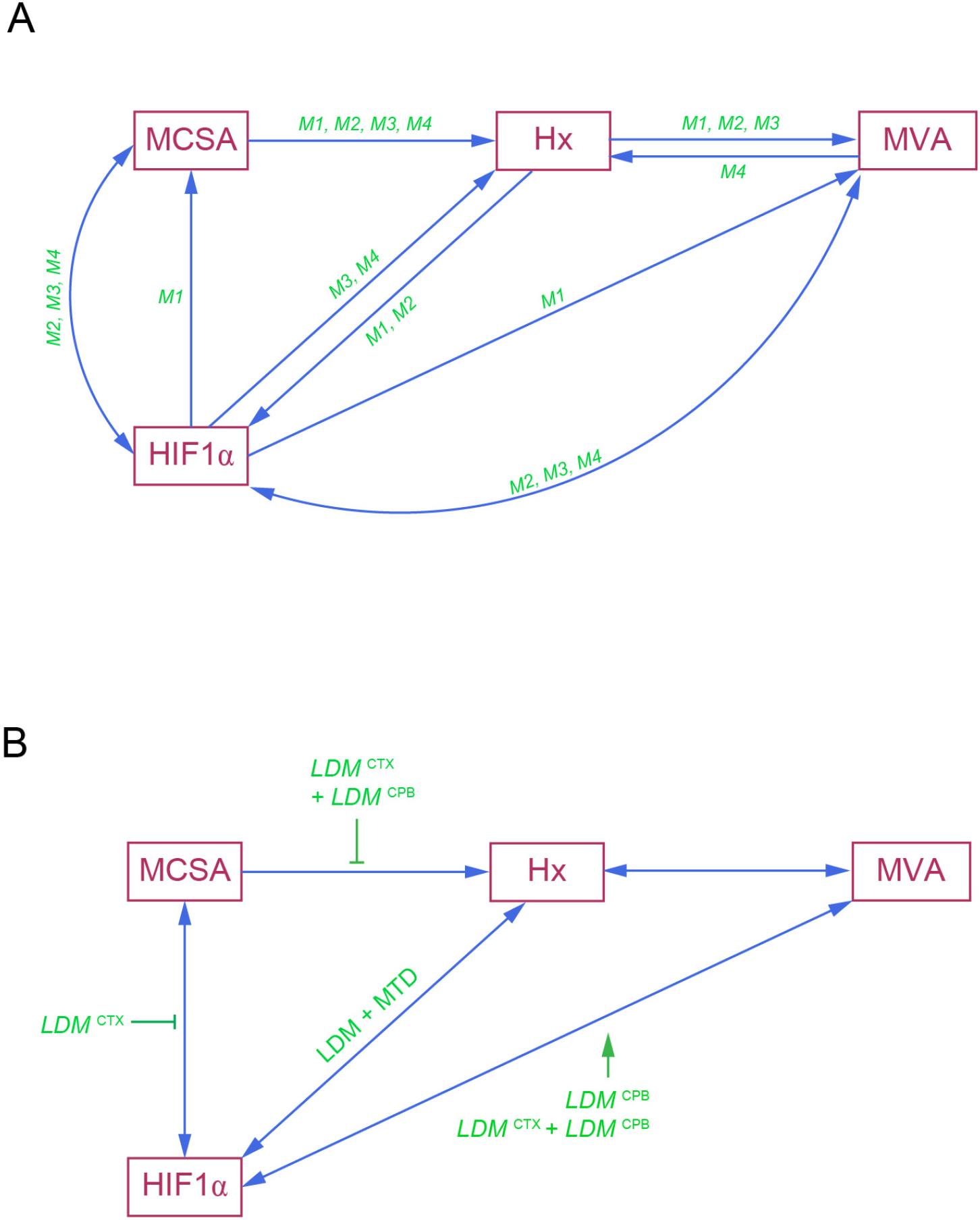
Integration of final path models and chemotherapy modality effects. **A**, Diagram showing all interactions among nodule size, hypoxia, HIF-1α and vascularisation in models M1 through M4. **B**, Effects of singlet/doublet low-dose metronomic and conventional, maximum-tolerated dose chemotherapy regimens upon *consensus* paths and covariances. *CTX*, cyclophosphamide; *CPB*, capecitabine; *HIF-1α*, hypoxia-inducible factor-1α; *Hx*, hypoxia; *LDM*, low-dose metronomic chemotherapy; *MTD*, maximum-tolerated dose chemotherapy; *MCSA*, metastatic cross-sectional area; *MVA*, metastatic vascular area. Double arrows indicate covariances, whereas single arrows indicate unidirectional paths. Individual treatments and their effects are depicted in *green*, wherein normal arrows show significant standardized loadings for a given path/treatment modality, whilst *broken arrows* indicate disruption (non-significance) of a specific path.

## Discussion

In this study, we performed a comprehensive path analysis of nested models correlating measures of metastatic size, intra-metastatic hypoxia, HIF-1α expression and vascular area utilizing a previously published dataset in an advanced model of breast cancer that disseminates to the lungs. The goal of this study was to deconvolve the complex relationships among these variables in order to uncover novel hypothetical pathways amenable for further experimental enquiry. To our knowledge, this is the first study utilizing path analysis to dissect hypoxic signalling in the metastatic tumour microenvironment whilst exploring the effects of different chemotherapeutic modalities upon intra-metastatic hypoxia.

Notwithstanding the wealth of evidence indicating that HIF-1α can promote every single step down the metastatic cascade (Schito & Semenza, 2016; Schito & Rey, 2017), the role of hypoxia and HIFα on metastatic tumours remains less well-defined (Cao *et al*, 2009; van der Wal *et al*, 2012; Shimomura *et al*, 2013). In this study, we sought to offer statistical insights into the interactions among metastatic tumour size, hypoxia, HIFα signalling and microvascular area *via* path analysis, a component of structural equation modelling that allowed us to take a combinatorial approach in order to compare hypothetical models, to potentially generate alternative hypotheses, and to test the robustness of established mechanisms driving hypoxic metastatic progression, in a model of advanced breast cancer (Schito *et al*, 2020). In our previous study, we observed that LDM chemotherapy was able to offset the well-described in vitro induction of HIF-1α by conventional MTD chemotherapy (Cao *et al*, 2013; Samanta *et al*, 2014). As a result of these data, we sought to evaluate whether the effect of LDM in metastatic breast tumours colonizing the lungs was due to inhibition of HIF-1α-driven vascularisation (angiogenesis) whilst examining the presence of HIFα independent effects.

Our analysis showed that the interactions amongst metastatic tumour size, hypoxia, HIFα signalling and microvascular area were convergent upon a general path organization wherein tumour growth induces diffusion-limiting intra-tumoral hypoxia, which in turn is strongly associated with an increase in microvascular area (angiogenic response). GLMs derived from the data revealed a significant statistical interaction between HIF-1α and hypoxia, whereas all other variables were orthogonal. Unexpectedly, HIF-1α levels acted as an intermediate (mediator) variable in-between metastatic tumour size and hypoxia, thus interacting with both, albeit suggesting direct, HIF-1α-independent effects controlling both tumour growth and intra-metastatic vascularisation and oxygenation. By contrast, the association between HIF-1α and vascularisation was less consistent across models; likewise, we observed that reversing the direction of the path between hypoxia and HIF-1α did not improve model goodness-of-fit. Collectively, these data suggest that HIF-1α signalling does not act as *a sine qua non* mediator of metastatic growth and angiogenesis; thereby raising the possibility of bi-directional relationships between hypoxia and HIF-1α levels.

Another salient finding from this study was the observation that hypoxia and HIF-1α both correlate in a positive, rather than in a counteracting manner within experimental metastatic breast tumours. This finding is at odds with the notion that an increase in tumour vascularisation *via* angiogenesis should improve perfusion and therefore oxygenation, thereby staving-off hypoxia. Alternatively, our data is in line with recent experimental data indicating that tumour angiogenesis generates dysfunctional vessels that paradoxically worsen perfusion and deepen hypoxia (Jain, 2014; Schito *et al*, 2020; Schito & Rey, 2020). From this perspective, increasing perfusion and oxygenation represents a desirable outcome in order to improve the access of targeted therapeutics upon cancer cells in angiogenic tumours, a concept known as vessel normalisation (Jain, 2005; Hamzah *et al*, 2008; Martin *et al*, 2019). Likewise, our modelling data support the notion of microvascular density as a correlate of intra-tumoral hypoxia and angiogenic activity, rather than an indicator of angiogenic activity, a concept originally noted by Judah Folkman (Hlatky *et al*, 2002). Furthermore, treatment with either LDM or MTD chemotherapy regimens disrupted the correlation between hypoxia and HIF-1α as opposed to LDM, which promoted the correlation between HIF-1α and vascularisation. In addition, singlet and doublet LDM regimens disrupted the correlation between metastatic size, hypoxia and HIF-1α without effect upon vascularisation, thus suggesting that the offsetting effect of LDM chemotherapy upon HIF-1α is likely to exert cell-autonomous effects (*e*.*g*., metabolism and O_2_ consumption), rather than direct antiangiogenic effects of relevance for personalised targeting of hypoxia in cancer (McIntyre & Harris, 2015).

Taken together, our findings provide novel working hypotheses by revealing hitherto unknown statistical interactions wherein metastatic tumour growth interacts with hypoxia, HIF-1α signalling and angiogenic responses, as well as suggesting a non-angiogenic, differential effect of both metronomic and maximum-tolerated dose chemotherapy regimens upon the hypoxic metastatic tumour microenvironment. This work highlights the potential of path analysis and structural equation modelling approaches as tools statistically model the hypoxic tumour microenvironment and hypothesize mechanisms of action for conventional and targeted oncolytic agents.

## Acknowledgements

LS and SR-K are supported by the UCD *Ad Astra* Fellows Programme and UCD School of Medicine, University College Dublin, Ireland.

## Author contributions

SR-K and LS conceived the study, analysed data, developed tools for quantitative immunohistochemistry and coding scripts for statistical analyses, drafted and wrote the manuscript. SR-K and LS reviewed and approved the contents of the manuscript.

## Data availability

The raw data analysed in this study are available upon request.

